# Structure and dynamics of RNA guanine quadruplexes in SARS-CoV-2 genome. Original strategies against emerging viruses

**DOI:** 10.1101/2021.05.20.444918

**Authors:** Tom Miclot, Cécilia Hognon, Emmanuelle Bignon, Alessio Terenzi, Marco Marazzi, Giampaolo Barone, Antonio Monari

**Author notes:** Electronic Supplementary Information (ESI) available: Protocol of the sequence and sequence-structure analysis, and additional details of the MD and QM/MM simulations RNA G4 structural parameters for the different replicas and unshifted ECD spectra. Pdb of RG-1 structure See DOI: 10.1039/x0xx00000x.

## Abstract

Guanine quadruplexes (G4) structures in viral genome have a key role in modulating viruses’ biological activity. While several DNA G4 structures have been experimentally resolved, RNA G4s are definitely less explored. We report the first calculated G4 structure of the RG-1 RNA sequence of SARS-CoV-2 genome, obtained by using a multiscale approach combining quantum and classical molecular modelling and corroborated by the excellent agreement between the corresponding calculated and experimental circular dichroism spectra. We prove the stability of RG-1 G4 arrangement as well as its interaction with G4 ligands potentially inhibiting viral protein translation.

---

At the end of 2019, a sudden outbreak of Severe Acute Respiratory Syndrome (SARS) developed in mainland China,^1^ to further spread worldwide obliging the World Health Organisation (WHO) to declare the emergence of a pandemic in March 2020. The syndrome is caused by a novel coronavirus, SARS-CoV-2, and has been styled COVID-19.^2,3^ Despite the relatively low mortality, SARS-CoV-2 is highly contagious and COVID-19 can evolve into severe forms necessitating critical care. Hence, COVID-19 is causing a considerable strain on healthcare systems, requiring unprecedented large-scale social distancing and containment measures, including full lockdowns. Even though vaccines have been promptly developed and released,^4^ also including the emergent messenger RNA (mRNA) technology,^5^ COVID-19 is still raging Worldwide in middle Spring 2021. The harmful effects of COVID-19 are also aggravated by the fact that no clearly efficient and safe antiviral agent has been proposed for large scale use. Indeed, while some nucleoside analogues, including Remdesivir, have shown high antiviral efficiency *in vitro* and *in vivo*, related side-effects strongly hamper their diffusion.^6^ In parallel to the mobilization of the medicinal chemistry community,^7^ several structural biology^8^ as well as molecular modelling and simulation^9^ groups have produced an unprecedented effort, which has resulted in the resolution and characterization of the main SARS-CoV-2 structural and non-structural proteins.

The rather complex organization of viral genome, also in case of RNA viruses, has been recently highlighted and related to their biological functions. In particular, the presence of guanine-quadruplexes (G4) arrangements has been spotlighted.^10^ The presence of quadruplexes folding may preserve the viral genetic material avoiding its recognition by the immune system. On the other hand, it has been shown that an overstabilization of G4s may inhibit the translation of viral proteins by the cellular apparatus. In the case of SARS-type coronaviruses it has also been shown that the highly-conserved SARS Unique Domain (SUD), used to sequestrate pro-apoptotic cellular mRNA sequences, is maintained in its active dimeric form by the interaction with G4 RNA sequences.^11–13^ As a matter of fact, while SARS-CoV-2 genome is in some extent less prone to arrange in quadruplexes, compared to other viruses such as Zika,^14^ four putative G4 sequences have been recently evidenced by Zhao *et al.*^15^ In particular the so-called RG-1 sequence, located in the nucleocapsid (N) protein coding region, has been characterized using electronic circular dichroism (ECD). The presence of G4s in infected living cells has also been confirmed, and their stabilization by ligands can induce the downregulation of their expression, impairing the maturation and infectivity of viral proteins hence paving the way to appealing therapeutic strategies.

Despite the importance of RNA G4 in the biological cycle of viruses like SARS-CoV-2, their structural characterization remains usually elusive. This is also underlined by the relative scarce number of RNA G4 structures that have been resolved, especially compared to their DNA counterparts. At the same time, the high ECD sensitivity to secondary structure rearrangements allows to achieve a molecular resolution giving access to all the subtle structural factors that may be crucial in driving the possible interactions with external ligands. In this communication we study the RG-1 sequence through a combination of multiple sequence alignment, homology modelling, classical molecular dynamics (MD), and hybrid quantum mechanics/molecular mechanics (QM/MM), in order to disentangle all the structural factors associated to the G4 conformations, also by comparing the Time Dependent Density Functional Theory (TD-DFT) simulated ECD spectrum with the experimental one by Zhao *et al*.^15^ The full computational strategy of our multiscale approach can be found in Electronic Supplementary Information (ESI).

The structure of RG-1 was predicted from a multiple sequence alignment with three DNA G4s sequences corresponding to experimental structures (PDB codes 2N4Y,^16^ 6T51,^17^ 5I2V).^18^ The RG-1 structural model exhibits a parallel G4 composed of two superposed tetrads – see Figure 1. Interestingly, we can also evidence that, in addition to the G4 core, a rather large non-structured loop, composed of adenines and cytosines, is also present as a linker to the guanines involved in the tetrad. The same loop is also present on the human DNA G4 used as template (PDB 6T51) – see Figure 1-B. Two independent 1 μs MD trajectories of the RG-1 G4 model solvated in a periodic box of TIP3P water^19^ have been carried out using the amber f99 force field including bsc1 corrections^20,21^ with the NAMD code.^22^ For sake of clarity, the results of the first simulations are described below and the description of the second simulation, which exhibit identical trends, can be found in ESI.

**Figure 1.**
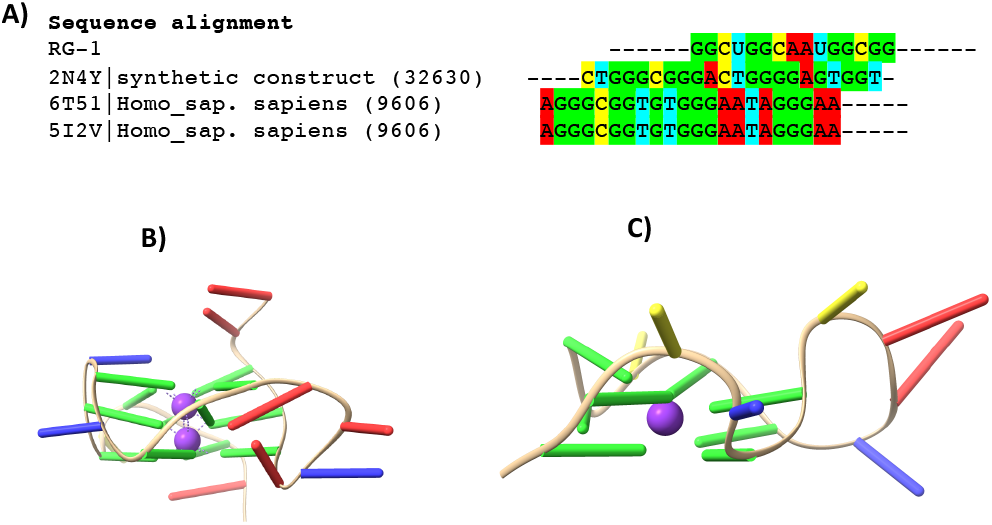
A) Alignment of RG-1 sequence with a synthetic construct from HIV-1 genome (2N4Y) and two homo sapiens DNA G4s sequences (6T51 and 5I2V). B) Side view of the structure of the G4 DNA from PDB 6T51 and C) of the reconstructed RG-1.

From our MD simulations, the RG-1 tetrad core is extremely stable and only experiences slight vibrational deformation – see Figure 2-A. As expected, the connecting loop is much more flexible and accesses a larger conformational space. Interestingly, as reported in ESI, the fluctuations of this loop also lead to some metastable states in which adenines and cytosines can be π-stacked to the guanines in the tetrad.

**Figure 2.**
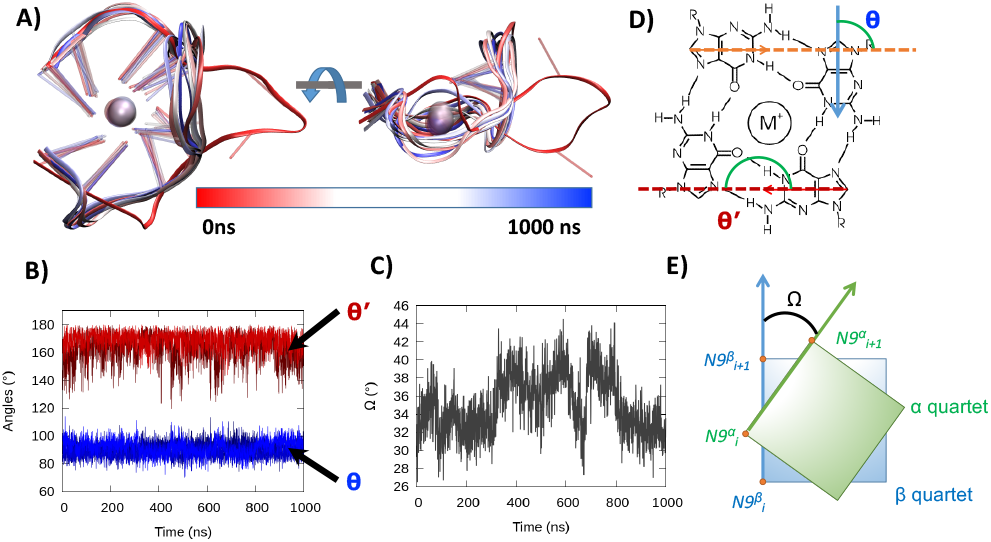
A) Representative snapshots of the RG-1 RNA G4 extracted along the MD trajectory and superposed, the color code represents the time evolution. Time evolution of the θ and θ’ angles for the first, α, tetrad (B) and of the twist angle Ω (C) Schematic depiction of θ and θ’ angles indicating the arrangement of guanines on the tetrad (D) and of the twist angle Ω (E).

Core rigidity and loop flexibility can also be inferred from the analysis of the Root Mean Square Deviation (RMSD) - see ESI. While the RG-1 sequence presents a moderate RMSD peak at around 8.0 Å, this value is mainly due to the contribution of the peripheral loop, which amounts for 6.5 Å. On the contrary, the RMSD of the nucleobases in the tetrads barely reaches 2.6 Å, highlighting the negligible deviation of the tetrad core from the original starting conformation.

As usual in G4 arrangements, the rigidity of the central core is due to the involvement of the guanines in the Hoogsteen hydrogen bonding network and in the favourable interaction with the metal cations present in the central channel.

The stability of G4 arrangement is reflected in the almost ideal values assumed by the θ and θ’ angles defining the disposition of the guanines in the tetrad – see Figure 2-D. θ peaks at 90.4 ± 4.6°, and θ’ is centred around 165.7 ± 8.8° in both tetrads – see Figure 2-B and ESI. Such a situation is indicative of the formation of a stable and persistent Hoogsteen hydrogen bonding network, hence confirming the propensity of the RG-1 RNA sequence to assume a G4 parallel conformation. As a more global descriptor, the stability and rigidity of the G4 core is also reflected by the time evolution of the Ω angle describing the twisting between the two tetrads (Figure 2-C and 2-E), which is centred around the ideal value of 33.1 ± 2.2° typical of a parallel arrangement. The stability of Ω is also more evident in the case of the second trajectory (32.3 ± 2.3 see ESI) in which hardly any deviation from the ideal value can be observed.

Our MD trajectories consistently and unequivocally confirm that the RG-1 sequence of the SARS-CoV-2 genome is able to assume a stable parallel G4 conformation. In their original work, Zhao *et al*.^15^ have used ECD spectroscopic signatures to confirm the structuration of the RNA sequence into a G4 arrangement. To allow a one-to-one mapping between molecular modelling and experimental results we have simulated the ECD spectrum using a hybrid QM/MM approach on top of snapshots extracted from the MD trajectory, following a protocol successfully used by our group for related systems.^23–25^ The QM/MM excited state calculations have been performed using the Orca/Amber interface, and the eight nucleobases forming the G4 core have been included in the QM partition to consider the coupling between the π-stacked chromophores (Figure 3-B). The results are in very good agreement with the experimental spectrum published by Zhao *et al*. ^15^ – see Figure 3. The experimental spectrum presents a first large and rather intense positive peak centred around 270 nm followed by a negative peak at 240 nm, whose intensity is much smaller, the ratio being about 1:4. Such a pattern is common in the case of nucleic acid aggregates and can be seen as a typical spectroscopic feature of parallel G4 arrangement. The results of our QM/MM simulation produced an ECD spectrum with similar pattern, but with blue-shifted signals, due to the use of a reduced-size basis set.

**Figure 3.**
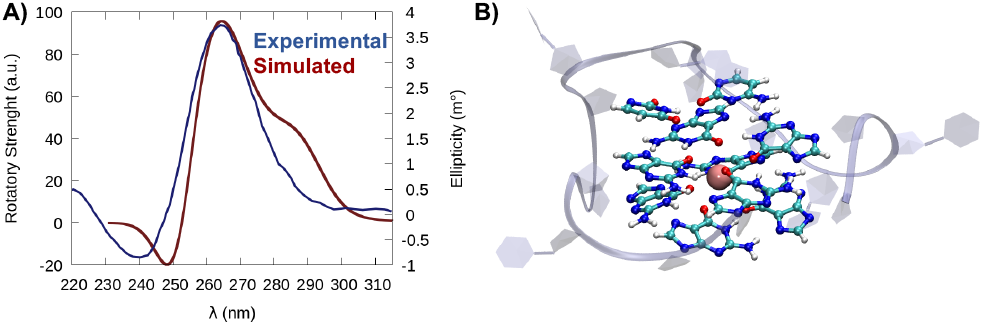
A) Experimental, from Zhao et al,^15^ and simulated ECD spectrum of the RG-1 RNA sequence. Note that the simulated spectrum has been homogeneously shifted by 40 nm. The simulated spectrum is obtained by QM/MM at TD-DFT level with M06-2X functional and the 6-31G(d) basis. The chosen QM partition is highlighted in ball and stick representation while the G4 backbone and dangling bases are shown in cartoon (B).

Although a larger basis set could reproduce with more precision the absorption energy, in the context of the present work we considered of more prominence the correct description of the global shape of the ECD signal, thus confirming the G4 conformation, at the same time correctly describing the electronic nature of the spectral signature. The choice of the basis set was imposed not only by the computational overload due to the extended QM partition, but also by the necessity to avoid QM wavefunction overpolarization due to the interaction with the MM point charges. To palliate to the basis set incompleteness, we apply a global shift of 40 nm to the simulated ECD signal to allow the straightforward comparison with the experimental counterpart – see Figure 3. The original non-shifted ECD spectrum is reported in ESI. The similarity between the simulated and experimental signals are self-evident, providing a clear picture in terms of the relative position of the two main peaks, their intensity, and the global band shape: a broad positive band is followed by a less intense, and slightly sharper, negative signal. The excellent agreement between the calculated and experimental spectra provides a further definitive confirmation of the folding of the RG-1 sequence in a parallel G4 conformation.

Having shown the stability and the persistence of the G4 arrangements, the question arises whether some ligands could interact with the RG-1 RNA sequence, and possibly overstabilize it to induce an effective inhibition of the translation of the viral genome and impair the viral cycle. In this context, we have examined the interaction of two ligands with the RG-1 sequence, namely quarfloxin (QRF) and pyridostatin (PDP). QRF (also called CX-3543) is a well-known G4-binding ligand, believed to target RNA G4s and evaluated in phase II clinical trials for human cancer therapy.^26^ PDP is also known for its capacity to modulate the expression of G4 containing genes, providing antiproliferative effects.^27^ Interestingly, PDP was reported by Zhao *et al*.^15^ as a ligand capable of selectively recognize RG-1 and increase its melting temperature. The initial complexes between RG-1 and the two ligands have been obtained by docking the drugs onto the parallel G4 structure. The stability of each significant pose has been further confirmed running two independent 1 μs MD trajectories. Representative structures of the main binding modes are reported in Figure 4, and the results for a slightly different initial pose, in which the ligand interacts with the tetrad are also collected in ESI.

**Figure 4.**
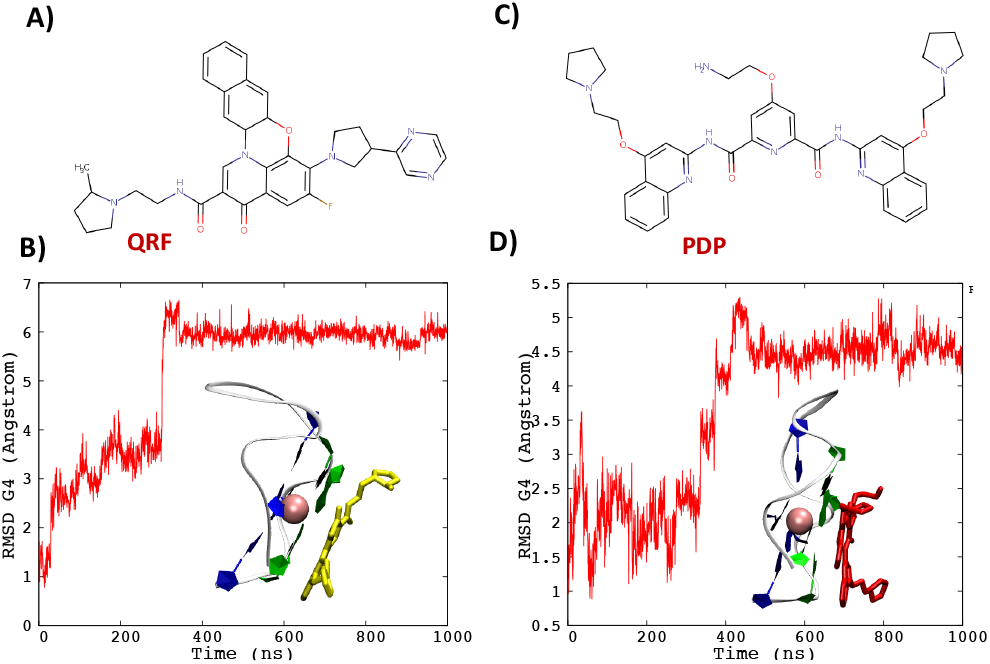
Chemical representation of quarfloxin, QRF (A) and pyridostatin, PDP (C) together with a representative snapshot of the ligand binding mode and the time evolution of the RMSD (B,D respectively).

Both the docking and the MD simulations agree in previewing the formation of stable aggregates between the RNA G4 and the small ligands, the persistence and stability of which can also be appreciated by the extended plateau observed on the RMSD time-series after 400 ns (Figure 4-B and D). More importantly, it can be observed that the binding takes place via π-stacking of the ligand on top of the tetrad plans, and it is mostly driven by dispersion and hydrophobic interactions. This result can be rationalized due to the presence of conjugated moieties on the ligands, and their globally planar structure. Furthermore, such interaction modes, while providing an enhanced stabilization of the G4 arrangement due to the increase of the attractive interactions, induce only moderate or negligible structural perturbation as can also be observed from the angles between the guanines and the twist reported in ESI for the different interaction modes and ligands. The structural stability of the G4 arrangement upon binding with ligands is also consistent with the experimental results that showed only slight differences in the ECD spectra upon interaction of RG-1 with PDP. In addition, the π-stacking interaction, also due the largely aromatic nature of both ligands, is susceptible to take place without any important energetic barrier, or steric hindrance, hence greatly facilitating the recruitment of the drug. While incontestably pointing towards the formation of stable aggregates, the results of the equilibrium MD simulations alone cannot underly any significant or qualitative difference between QRF and PDP binding, hence suggesting that both can potentially be seen as valuable ligands to stabilizing G4 arrangements and more particularly the parallel conformation of the RG-1 RNA sequence. We have unravelled the structural behaviour of a putative G4 RNA sequence present in the genome of SARS-CoV-2 using multi-scale molecular modelling approaches. The combination of the multiple sequence alignment, the μs-scale sampling of the conformational space and computational spectroscopy clearly demonstrated that the RG-1 sequence adopts a stable parallel G4 conformation composed of two rigid tetrads and a flexible peripheral loop. In addition, we have shown that the G4 conformation can, without any major structural rearrangement, form stable aggregates with known G4 ligands, that are susceptible to increase the persistence of the quadruplex structure. The important role of G4s in tuning the viral response and the biological cycle, including emerging RNA viruses such as Zika, Dengue, or coronaviruses, calls for the precise determination of putative G4 sequences. Moreover, influencing the equilibrium between unfolded and G4 sequences via the use of small drugs offers an original, yet not fully explored, possibility for the development of novel and potentially wide-action antiviral agents. This study highlights the robustness of our *in silico* protocol to provide a most favourable complement to experimental studies in suggesting specific interaction modes and structures of RNA sequences. Our contribution represents a proof of concept of the capacities offered by mature and multiscale simulation techniques to unravel key biological processes and phenomena.

The work has been conducted via the financial support of the Universities of Palermo and Lorraine, and the French CNRS. T.M. thanks the University of Palermo for granting a Ph.D. fellowships. Calculations have been partially performed on the local LPCT computing cluster and on the Regional Explor Computing Center. The authors thank GENCI for providing access to the national computing center under the project “Seek&Destroy”. M.M. is grateful to the University of Alcalá for providing funds under the COVID-19 project 2020/00256/001. There are no conflicts to declare. E.B. and A.M. thank the French Research Ministry (MESRI) for funding under the GAVO project.

## Supporting information

Electronic Supplementary Inforlmation

## Notes

### Competing Interest Statement

The authors have declared no competing interest.

