## Supplementary material for "Structure and dynamics of RNA guanine quadruplexes in SARS-CoV-2 genome. Original strategies against emerging viruses": Electronic Supplementary Inforlmation

**Electronic Supplementary Information for**

*a. Department of Biological, Chemical and Pharmaceutical Sciences. University of Palermo, via delle Scienze 90126 Palermo, Italy.*

*b. Université de Lorraine and CNRS, LPCT UMR 7019, F-54000 Nancy, France.*

*c. Departamento de Química Analítica, Química Física e Ingeniería Química, Universidad de Alcalá, Ctra. Madrid-Barcelona Km. 33,600 E-28805, Alcalá de Henares (Madrid), Spain*

*d. Instituto de Investigación Química "Andrés M. del Río" (IQAR), Universidad de Alcalá, Ctra. Madrid-Barcelona Km. 33,600 E-28871, Alcalá de Henares (Madrid), Spain*

*

### Protocol for the set-up of a suitable initial geometry of the RG-1 RNA sequence in G4 arrangement.

A) Search for homologous G4 structures.

The RG-1, present in SARS-COV-2 genome, has been highlighted as a putative G4-forming RNA sequence, however no structural resolution has been attempted, hence we started by looking to homologous G4 sequences. The number of resolved RNA G4 structure are much lower than their DNA counterpart, so we considered also DNA arrangements considering thymine an analogue of uracil. Furthermore, considering that the experimental ECD spectrum of the RG-1 G4, reported by Zhao et al.,[1] we restricted our search to sequences folding in a parallel conformation.

The structures present in the RCSB PDB database have been mined imposing two criteria: sequence homology with the RG-1 target, and structural homology with the h-telo G4 DNA structure (PDB:1KF1) used as a template to force parallel G4 arrangement. It has to be noted that since only 15 residues are present in the RG-1 sequence we have been obliged to resort to motif search instead of full sequence search. The relaxed structure search and PROSITE[2] have been used.

While a first search on RNA structure target did not return any structure, enlarging the database to include DNA structure yielded 108 matches, of which only the first five high ranking structures have been retained since most of the others were actually referring to double helix B-DNA (Table S1).

| PDB Code | Nucleic type | Organism | Sequence |
|---|---|---|---|
| 2N4Y | DNA | Synthetic construct (32630) | CTGGGCGGGACTGGGGAGTGGT |
| 6T51 | DNA | *Homo sapiens sapiens* (9606) | AGGGCGGTGTGGGAATAGGGAA |
| 5DWX | DNA | Synthetic construct (32630) | GGGTGGGTGGGTGGGTTAGCGTTA (chain A) |
| 5I2V | DNA | *Homo sapiens sapiens* (9606) | AGGGCGGTGTGGGAATAGGGAA |

**Table S1:** DNA structures sorted by the RSC PDB data base scan and forming parallel G4 arrangements.

B) Refinement of the selected G4 structures.

Firstly, Clustal Omega software[3] was used to perform a sequence alignment between the RG-1 sequence and the five structures reported in Table S1, as shown in Figure S1-A. Note that to match the sequences obtained from the PDB mining, RG-1 was manually and artificially transformed in a DNA sequence.

All the selected DNA G4 structures are constituted by the alignment of 3 tetrads, whereas the GR-1 RNA features only guanine plans. This can be easily inferred since all the guanines are organized in groups of three, separated by other nucleotides. It will therefore be necessary to take this aspect into account during the RG-1 RNA reconstruction process. Secondly, the alignment shows that while the PDB:5DWX structure does indeed show some sequence similarities with our target, it misses a crucial structural feature, i.e. an extended loop bridging the tetrads that is instead evident in RG-1, hence this particular sequence has not been considered anymore. On the contrary, all the other sequences present the extended loop and hence share stronger similarity with our putative G4 forming sequence (Figure S1-B).

Considering the retained structures, PDB:6T51 and PDB:5I2V, both share the same sequence and correspond to the same DNA promoter, hence there should be no major difference between them. On the contrary, PDB:2N4Y presents an additional difficulty since it is constituted by 13 guanines. Thus, the additional guanine should not participate in the formation of the tetrads, and it can be positioned either in the large or small loop, or simply hanging at the border of the arrangement. Hence, to take into account all these aspects, we aligned the three structures following a structure-sequence protocol implemented in the UCSF Chimera software[4] (Figure S1-C). The robustness of the alignment and the similarities between the three structures can be appreciated from Figure S1, furthermore, it is also confirmed that the additional guanine (G13) of the PDB:2N4Y structure does not participate in the formation of the tetrads, but it is instead located in the extended loop. Considering the similarity of the structures, the absence of additional guanines that could perturb the G4 dynamics, and the fact that differently from the other two PDB:2N4Y does not correspond to a biological relevant structure but is instead a synthetic construct, we chose PDB:5I2V as our template for RG-1.

C) Construction of the starting model of GR-1 RNA

The chosen PDB:5I2V was modified with Maestro software[5] to match a two tetrad G-quadruplex followed by a complete minimization to remove artificial constrains. Afterwards, the "Mutate" tool of COOT software[6] was used to edit the sequence and obtain the final RG-1 RNA by mutating the template residues, i.e. replacing thymine with uracil and deoxyribose with ribose.

**A) Sequence alignment**

```
RG-1|RNA converted to DNA                          -------GGCTGGCAATGGCGG------
2N4Y|synthetic construct (32630)                   ----CTGGGCGGGACTGGGGAGTGGT-
6T51|KRAS22RT promoter|Homo_sap. sapiens (9606)    AGGGCGGTGTGGGAATAGGGAA-----
5I2V|KRAS promoter|Homo_sap. sapiens (9606)        AGGGCGGTGTGGGAATAGGGAA-----
5DWX|Chain A|synthetic construct (32630)           -GGGTGG-GTGGG--TGGGTTAGCGTTA
5DWX|Chain B|synthetic construct (32630)           --------------------TAACGCTA
```

**B) Loop alignment**

```
RG-1|RNA converted to DNA                          --------GGCTGGCAATGGCGG-----
2N4Y|synthetic construct (32630)                   ----CTGGGCGGGACTGGGGAGTGGT-
6T51|KRAS22RT promoter|Homo_sap. sapiens (9606)    AGGGCGGTGTGGGAATAGGGAA-----
5I2V|KRAS promoter|Homo_sap. sapiens (9606)        AGGGCGGTGTGGGAATAGGGAA-----
```

**(C) Structure-Sequence alignment**

```
2N4Y|synthetic construct (32630)                   -----CTGG-GCGGGACT-G-GGG-AGTGGT--
6T51|KRAS22RT promoter|Homo_sap. sapiens (9606)    AGGGC--GGTGTGGGA-ATA-GGGA-------A
5I2V|KRAS promoter|Homo_sap. sapiens (9606)        AGGGC--GGTGTGGGA-A-TAGGGA------A-
                                                                  Loop
```

**Figure S1: (A)** Complete sequence alignment. Each base is represented with a specific color (green for guanine, yellow for cytosine, red for adenine, and light blue for thymine). Note that uracil in RG-1 has been replaced with thymine. **(B)** Alignment of the nucleobases bridging the guanine regions identifying the presence of an extended loop highlighted in purple **(C)** UCSF Chimera structure-sequence alignment of selected PDB structures.

### Molecular dynamic and QM/MM simulation protocol

A consistent protocol was used for all the classical MD simulations performed during this work.

All the initial structures were solvated in an octahedral TIP3P[7] water box with a buffer of 12 Å using Tlepa [2], negative charges were neutralized by adding $K^+$ ions to assure electroneutrality. Note that the central $K^+$ necessary for the G4 stability and inferred during the homology model was conserved (Figure S2). RNA was described by the amber ff99 force field[8] including bsc1 corrections.[9] MD simulations have been run on two independent replicas using the NAMD software,[10] with a hydrogen mass repartitioning (HMR),[11] which combined with the Rattle and Shake algorithms[12] allowed the use of a 4 fs time step. Each solvated system was minimized by performing 1000 steps of conjugated gradient followed by a total of 36 ns of equilibration gradually removing harmonic constrained on the RNA heavy atoms, followed by 1 µs of production for each replica. MD simulations have been performed in the isothermal and isobar (NPT) ensemble considering a temperature of 300 K and a pressure of 1 atm, maintained by Langevin isotherm thermostat[13] and piston,[14] Particle Mesh Ewald (PME)[15] was used consistently to treat long range electrostatic interactions with a cut off at 9 Å. The simulations were visualized and analyzed using VMD software[16] and G4 structural parameters have been extracted with the scripts provided by Tsvetkov *et al.*[17]

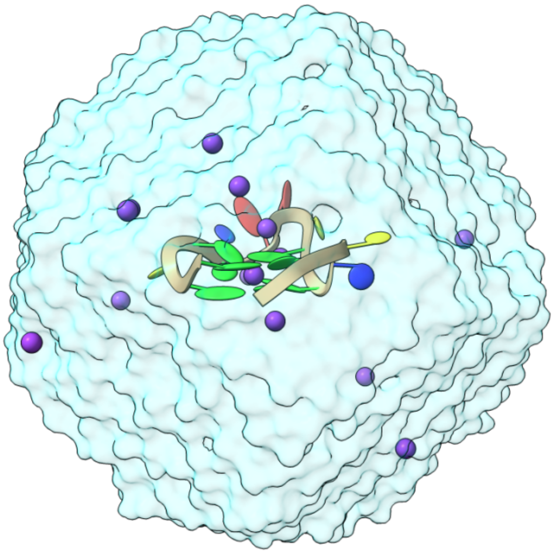

**Figure S2:** Representation RG-1 RNA solvated in an octahedral box of water and in the presence of $K^+$ ions.

### Molecular docking protocol

To build a suitable initial pose for the G4 bound to the ligand, molecular docking was used. A pdb file was generated from the last frame of the RG-1 MD simulation, to represent the receptor. Afterwards, the interaction between the RNA receptor and the ligands, i.e. quarfloxin and pyridostatin, was assessed by flexible docking with AutoDock Vina.[18] Two main poses, having quasi-degenerate score, singled out for both ligands corresponding to $\pi$-stacking between the ligand and the quartets, either on top (Figure S1-a) or on the bottom (Figure S1-b) Since the two poses are almost equivalent only one was deeply analyzed with subsequent MD simulation. Considering that both ligands are rather large and structurally rigid molecules, the interaction with the RNA backbone and grooves did not provide any stable conformation.

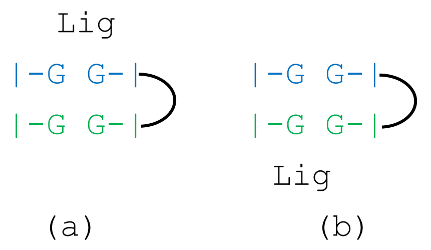


**Figure S3:** Schematic representation of the two best scoring docking pose involving either the interaction with quartet 1 (a) or quartet 2 (b). The large flexible loop is represented with the black circle segment.

### Parameterization of the ligand force field

The force field for the ligands were parameterized using the generalized amber force field (GAFF)[19] approach. The structure of each of the ligand was optimized at density functional theory (DFT) level using the B3LYP[20] functional and the 6-311+G(d,p) basis set. Atomic charges were obtained interpolating the restricted electrostatic potential (RESP)[21] obtained from an Harthree Fock calculation on top of the equilibrium geometry using the 6-31G(d) basis set, all the QM calculations have been performed using gaussian09.[22] The force field in its final form was obtained using the amber antechamber software.[23]

MD simulation in presence of the ligand was performed following the same protocol as the one described for solvated RG-1.

### Simulation of the ECD Spectra

ECD spectra have been simulated via a hybrid QM/MM approach obtaining vertical transitions on top of 100 snapshots randomly extracted from the MD simulation. The QM partition consisted of all the height guanines forming the G4 tetrads, the QM/MM frontiers being treated with the link atom approach, electrostatic embedding approach was consistently used. Vertical transitions have been calculated at time-dependent DFT (TD-DFT) level considering B3LYP, ωB97XD,[24] and M06-2X[25] functionals. 100 vertical transitions have been calculated for each snapshot. Only the 6-31G basis set was used, the relatively small basis set was used to cope with computational overload, due to the large size of the QM partition, and to avoid wave function overpolarization due to the interaction with nearby point-charges. The final spectrum is obtained convoluting the vertical rotatory strength and excitation energies with Gaussian functions of Fixed Width at Half Maximum (FWHM) of 0.3 eV to better match experimental results. Orca software[26,27] was used for the TD-DFT calculations and the QM/MM environment assured via the Amber external QM/MM interface.[28]

As can be seen from Figure S4, ωB97XD and M06-2X provides coherent results, differently from B3LYP which yields unphysical band shapes most probably due to the overstabilization of charge transfer states plaguing hybrid functionals. The differences between ωB97XD and M06-2X are mostly due to the fact that the former is globally more red-shifted and in particular the high-intensity positive peak at shorter wavelengths can already be observed in the space spanned by the calculated vertical transitions (100). The observed blue-shift can be ascribed to the small size of the basis set chosen.

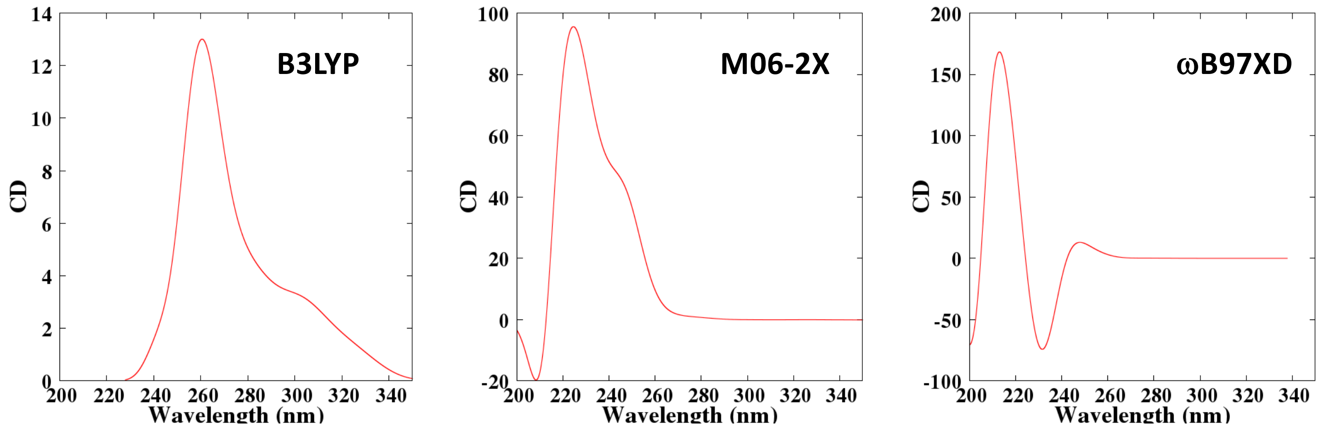


**Figure S4:** Simulated ECD spectra of the RG-1 RNA sequence obtained with different functionals on top of snapshots extracted from the MD simulation

22 M. J. Frisch, G. W. Trucks, H. B. Schlegel, G. E. Scuseria, M. A. Robb, J. R. Cheeseman, G. Scalmani, V. Barone, G.

A. Petersson, H. Nakatsuji, X. Li, M. Caricato, A. Marenich, J. Bloino, R. Janesko, R. Gomperts, B. Menucci, H. P. Hratchian, J. V. Ortiz, A. F. Izmaylov, D. Sonnenberg, F. Williams-Young, F. Ding, F. Lipparini, F. Edigi, J. Goings, B. Peng, A. Petrone, T. Henderson, D. Ranasinghe, V. G. Zakrzewski, J. Gao, N. Rega, W. Zheng, W. Liang, M. Hada, M. Ehara, K. Toyota, R. Fukuda, M. Hasegawa, T. Ishida, T. Nakajima, Y. Honda, O. Kitao, H. Nakai, T. Vreven, K. Throssell, M. J. J. A., J. E. Peralta, F. Ogliaro, M. Bearpark, J. J. Heyd, E. Brothers, K. N. Kudin, V. N. Staroverov, T. Keith, R. Kobayashi, J. Normand, K. Raghavachari, A. Rendell, J. C. Burant, S. S. Iyengar, J. Tomasi, M. Cossi, J. M. Millam, M. Klene, C. Adamo, R. Cammi, J. W. Ochterski, R. L. Martin, K. Morokuma, O. Farkas, J. B. Foresman, D. J. Fox, G. E. S. M. J. Frisch, G. W. Trucks, H. B. Schlegel, B. M. M. A. Robb, J. R. Cheeseman, G. Scalmani, V. Barone, H. P. H. G. A. Petersson, H. Nakatsuji, M. Caricato, X. Li, M. H. A. F. Izmaylov, J. Bloino, G. Zheng, J. L. Sonnenberg, T. N. M. Ehara, K. Toyota, R. Fukuda, J. Hasegawa, M. Ishida, J. Y. Honda, O. Kitao, H. Nakai, T. Vreven, J. A. Montgomery, E. B. J. E. Peralta, F. Ogliaro, M. Bearpark, J. J. Heyd, J. N. K. N. Kudin, V. N. Staroverov, T. Keith, R. Kobayashi, J. T. K. Raghavachari, A. Rendell, J. C. Burant, S. S. Iyengar, J. B. C. M. Cossi, N. Rega, J. M. Millam, M. Klene, J. E. Knox, R. E. S. V. Bakken, C. Adamo, J. Jaramillo, R. Gomperts, J. W. O. O. Yazyev, A. J. Austin, R. Cammi, C. Pomelli, G. A. V. R. L. Martin, K. Morokuma, V. G. Zakrzewski, A. D. D. P. Salvador, J. J. Dannenberg, S. Dapprich, and D. J. F. O. Farkas, J. B. Foresman, J. V. Ortiz, J. Cioslowski, G. M. J. Frisch, W. Trucks, H. B. Schlegel, G. E. Scuseria, M. A. Robb, J. R. Cheeseman, G. Scalmani, V. Barone, B. Mennucci, G. A. Petersson, H. Nakatsuji, M. Caricato, X. Li, H. P. Hratchian, A. F. Izmaylov, J. Bloino, G. Zheng, J. L. Sonnenberg, M. J. Frisch, G. W. Trucks, H. B. Schlegel, G. E. Scuseria, M. A. Robb, J. R. Cheeseman, G. Scalmani, V. Barone, B. Mennucci, G. A. Petersson, H. Nakatsuji, M. Caricato, X. Li, H. P. Hratchian, A. F. Izmaylov, J. Bloino, G. Zheng, J. L. Sonnenberg, M. Hada, M. Ehara, K. Toyota, R. Fukuda, J. Hasegawa, M. Ishida, T. Nakajima, Y. Honda, O. Kitao, H. Nakai, T. Vreven, J. A. Montgomery Jr., J. E. Peralta, F. Ogliaro, M. Bearpark, J. J. Heyd, E. Brothers, K. N. Kudin, V. N. Staroverov, R. Kobayashi, J. Normand, K. Raghavachari, A. Rendell, J. C. Burant, S. S. Iyengar, J. Tomasi, M. Cossi, N. Rega, J. M. Millam, M. Klene, J. E. Knox, J. B. Cross, V. Bakken, C. Adamo, J. Jaramillo, R. Gomperts, R. E. Stratmann, O. Yazyev, A. J. Austin, R. Cammi, C. Pomelli, J. W. Ochterski, R. L. Martin, K. Morokuma, V. G. Zakrzewski, G. A. Voth, P. Salvador, J. J. Dannenberg, S. Dapprich, A. D. Daniels, O. Farkas, J. B. Foresman, J. V. Ortiz, J. Cioslowski, D. J. Fox, W. C. Frisch, M. J.; Trucks, G. W.; Schlegel, H. B.; Scuseria, G. E.; Robb, M. A.; Cheeseman, J. R.; Scalmani, G.; Barone, V.; Mennucci, B.; Petersson, G. A.; Nakatsuji, H.; Caricato, M.; Li, X.; Hratchian, H. P.; Izmaylov, A. F.; Bloino, J.; Zheng, G.; Sonnenb, H. B. S. G. E. S. M. J. Frisch G. W. Trucks, G. S. V. B. B. M. M. A. Robb J. R. Cheeseman, M. C. X. L. H. P. H. G. A. Petersson H. Nakatsuji, G. Z. J. L. S. M. H. A. F. Izmaylov J. Bloino, R. F. J. H. M. I. T. N. M. Ehara K. Toyota, H. N. T. V. J. A. M. J. Y. Honda O. Kitao, M. B. J. J. H. E. B. J. E. Peralta F. Ogliaro, T. K. R. K. J. N. K. N. Kudin V. N. Staroverov, J. C. B. S. S. I. J. T. K. Raghavachari A. Rendell, J. M. M. M. K. J. E. K. J. B. C. M. Cossi N. Rega, J. J. R. G. R. E. S. V. Bakken C. Adamo, R. C. C. P. J. W. O. O. Yazyev A. J. Austin, V. G. Z. G. A. V. R. L. Martin K. Morokuma, S. D. A. D. D. P. Salvador J. J. Dannenberg, J. V. O. J. C. D. J. F. O. Farkas J. B. Foresman and D. J. Frisch, M. J.; Trucks, G.W.; Schlegel, H. B.; Scuseria, G. E.; Robb, M. A.; Cheeseman, J. R.; Scalmani, G.; Barone, V.;Mennucci, B.; Petersson, G. A.; Nakatsuji, H.; Caricato, M.; Li, X.; Hratchian, H. P.; Izmaylov, A. F.; Bloino, J.; Zheng, G.; Sonnenber, *Gaussian 09 Revis. D.01*, 2009, 2–3.

23 J. Wang, W. Wang, P. A. Kollman and D. A. Case, *J. Mol. Graph. Model.*, 2006, **25**, 247–260.

24 J. Da Chai and M. Head-Gordon, *Phys. Chem. Chem. Phys.*, 2008, **10**, 6615–6620.

25 Y. Zhao and D. G. Truhlar, *Theor. Chem. Acc.*, 2008, **120**, 215–241.

26 F. Neese, *Wiley Interdiscip. Rev. Comput. Mol. Sci.*, 2018, **8**, e1327.

27 F. Neese, *Wiley Interdiscip. Rev. Comput. Mol. Sci.*, 2012, **2**, 73–78.

28 A. W. Götz, M. A. Clark and R. C. Walker, *J. Comput. Chem.*, 2014, **35**, 95–108.

| | Run 1 | Run 2 |
|---|---|---|
| **RMSD (Å)** | 6.477 ± 0.598 | 6.618 ± 0.671 |
| **Tetrad Distance (Å)** | 3.470 ± 0.086 | 3.460 ± 0.091 |
| **Ω (°)** | 33.088 ± 2.291 | 32.368 ± 2.329 |
| **First Tetrad** | | |
| $\theta_{G1-G15}$ (°) | 91.425 ± 4.915 | 90.061 ± 4.786 |
| $\theta_{G1-G11}$ (°) | 89.908 ± 4.750 | 90.129 ± 5.108 |
| $\theta_{G5-G11}$ (°) | 88.128 ± 4.265 | 89.440 ± 4.346 |
| $\theta_{G5-G14}$ (°) | 92.146 ± 5.744 | 91.627 ± 4.861 |
| $\theta'_{G1-G14}$ (°) | 162.624 ± 10.598 | 168.232 ± 7.331 |
| $\theta'_{G11-G14}$ (°) | 168.975 ± 6.488 | 168.817 ± 7.017 |
| **Second Tetrad** | | |
| $\theta_{G2-G6}$ (°) | 89.792 ± 4.151 | 90.863 ± 4.191 |
| $\theta_{G2-G12}$ (°) | 89.790 ± 4.107 | 89.233 ± 4.417 |
| $\theta_{G6-G12}$ (°) | 92.661 ± 3.923 | 92.613 ± 4.386 |
| $\theta_{G6-G15}$ (°) | 89.758 ± 4.553 | 88.505 ± 4.511 |
| $\theta'_{G2-G15}$ (°) | 163.722 ± 9.223 | 165.129 ± 8.826 |
| $\theta'_{G12-G15}$ (°) | 166.529 ± 9.068 | 167.933 ± 8.412 |

**Table S2**: Average and standard deviation of the main structural parameters for the RG-1 simulations in the two replicas. The values of the θ and θ' angles are reported for each of the couple of guanines constituting the tetrads. The distance between the tetrads is considered as the distance between the center of mass of each quartet.

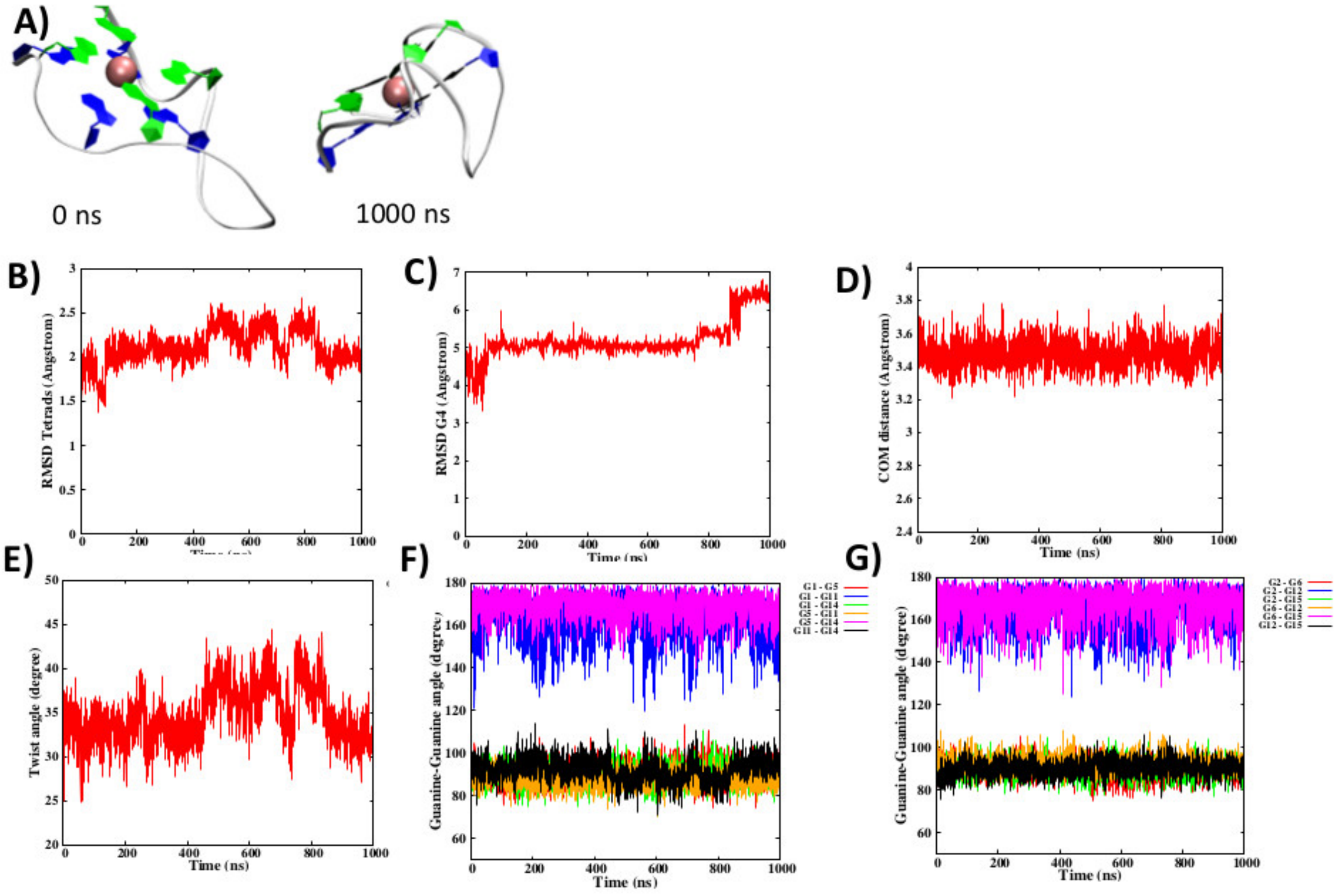


**Figure S5:** Initial and final snapshot of the first MD simulations of the RG-1 sequence (A). RMSD of the guanine in the tetrad (B) and of the flexible loop (C), distance between the tetrad (D), twist Ω angle (E), and ϕ, ϕ' angles for the first (F) and second (G) tetrad.

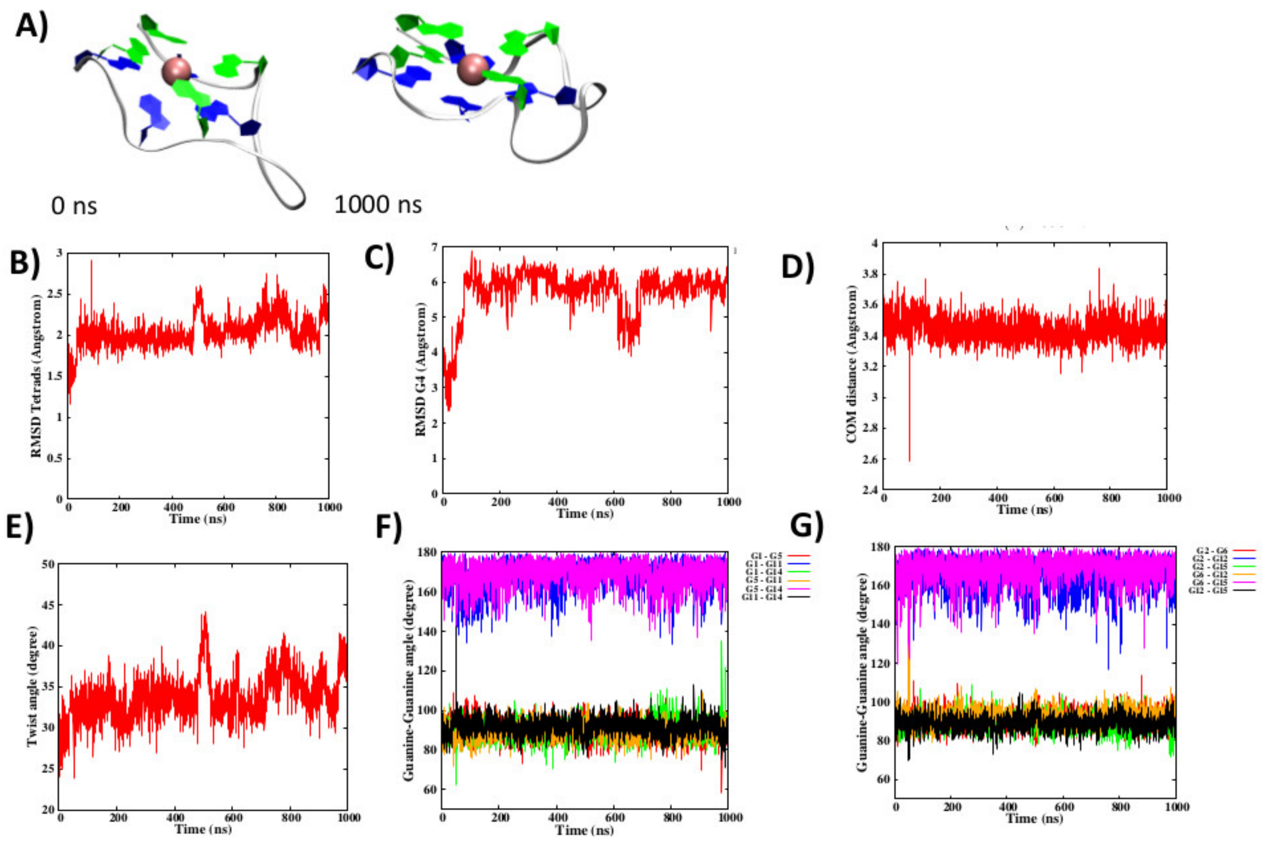


**Figure S6:** Initial and final snapshot of the second MD simulations of the RG-1 sequence (A). RMSD of the guanine in the tetrad (B) and of the flexible loop (C), distance between the tetrad (D), twist Ω angle (E), and ϕ, ϕ' angles for the first (F) and second (G) tetrad.

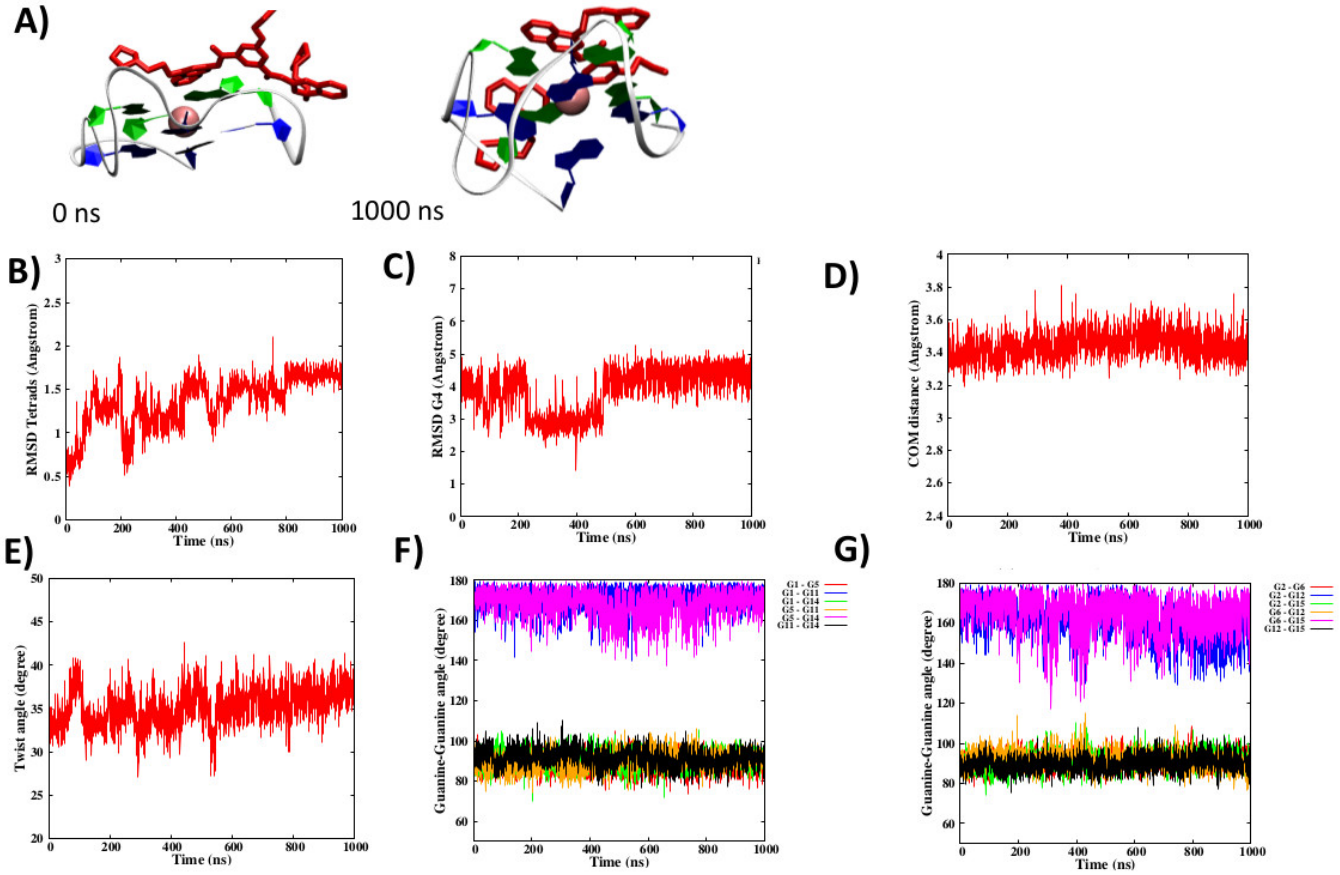


**Figure S7:** Initial and final snapshot of the first MD simulations of the RG-1 sequence interacting with PDP in the top binding pose (A). RMSD of the guanine in the tetrad (B) and of the flexible loop (C), distance between the tetrad (D), twist Ω angle (E), and ϕ, ϕ' angles for the first (F) and second (G) tetrad.

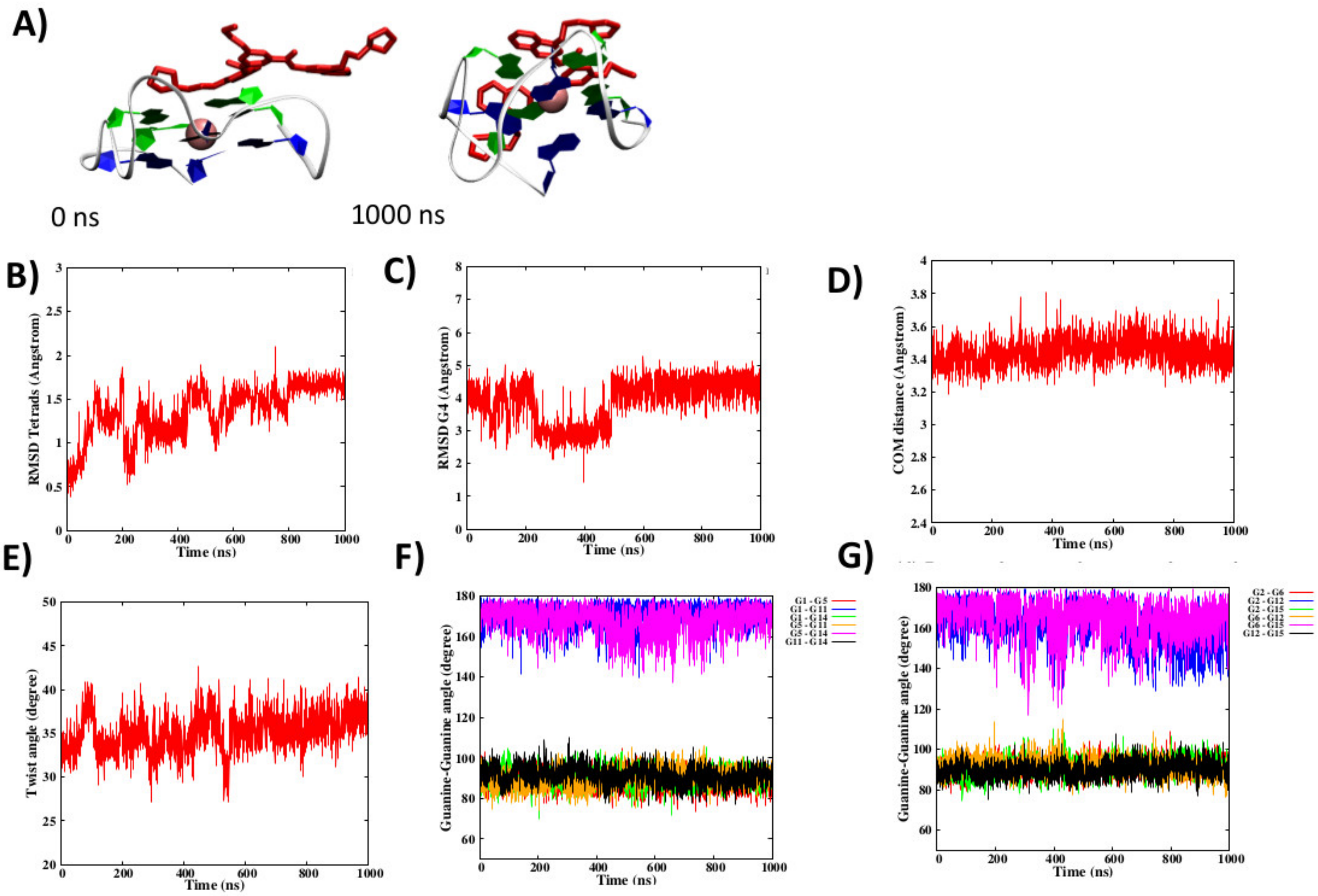


**Figure S8:** Initial and final snapshot of the second MD simulations of the RG-1 sequence interacting with PDP in the top binding pose (A). RMSD of the guanine in the tetrad (B) and of the flexible loop (C), distance between the tetrad (D), twist Ω angle (E), and ϕ, ϕ' angles for the first (F) and second (G) tetrad.

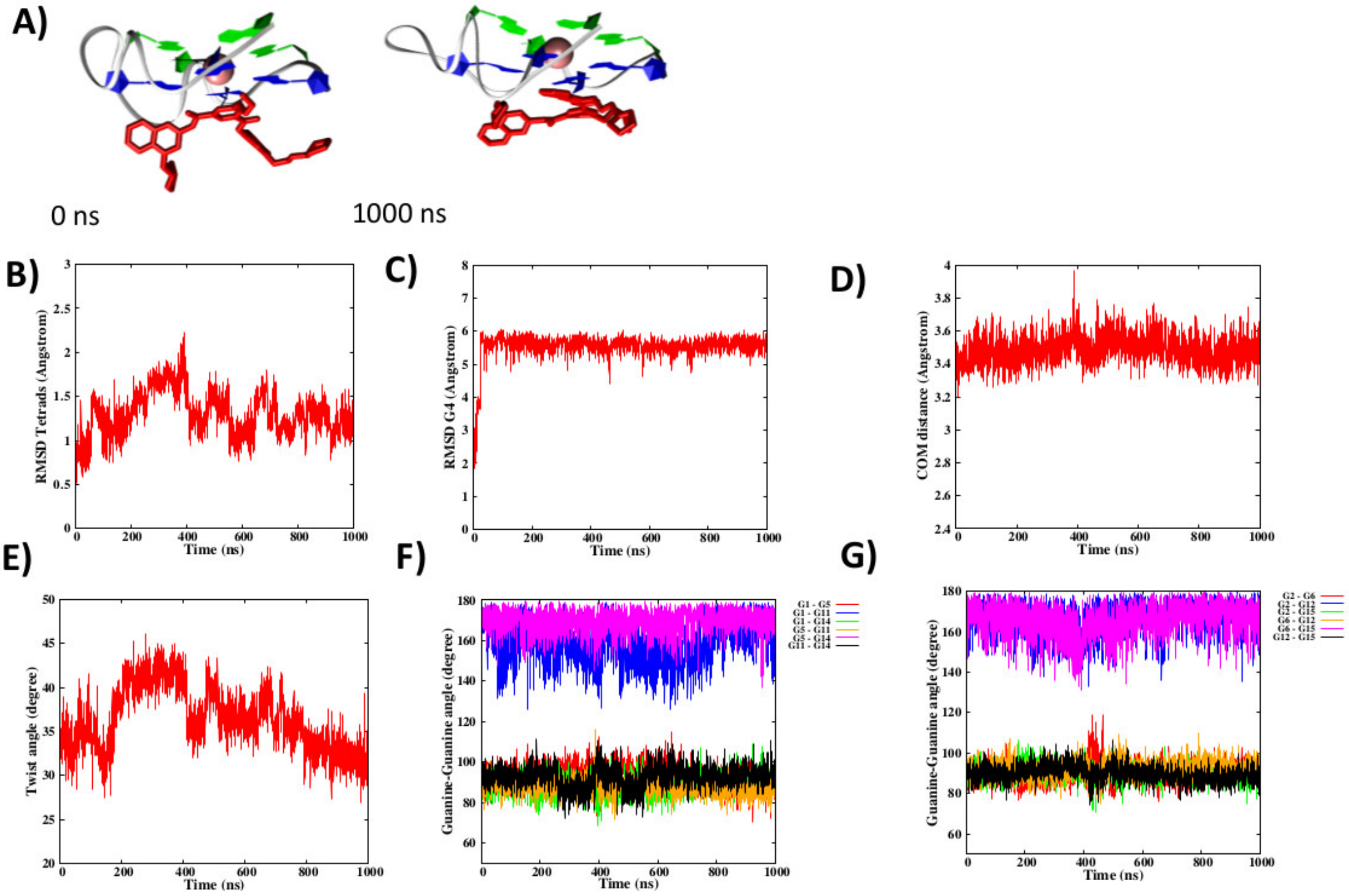


**Figure S9:** Initial and final snapshot of the first MD simulations of the RG-1 sequence interacting with PDP in the down binding pose (A). RMSD of the guanine in the tetrad (B) and of the flexible loop (C), distance between the tetrad (D), twist $\Omega$ angle (E), and $\phi$, $\phi'$ angles for the first (F) and second (G) tetrad.

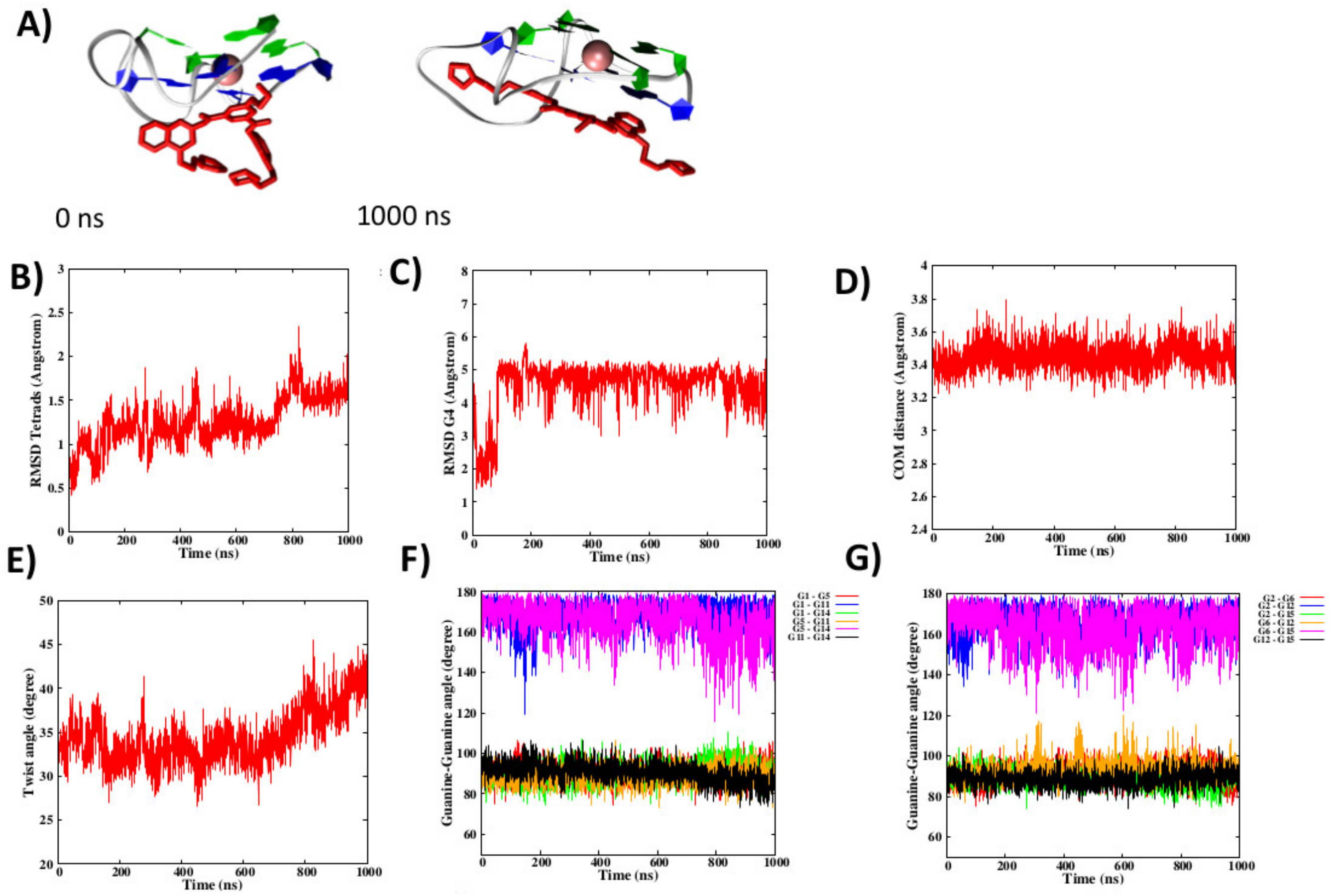


**Figure S10:** Initial and final snapshot of the second MD simulations of the RG-1 sequence interacting with PDP in the down binding pose (A). RMSD of the guanine in the tetrad (B) and of the flexible loop (C), distance between the tetrad (D), twist $\Omega$ angle (E), and $\phi$, $\phi'$ angles for the first (F) and second (G) tetrad.

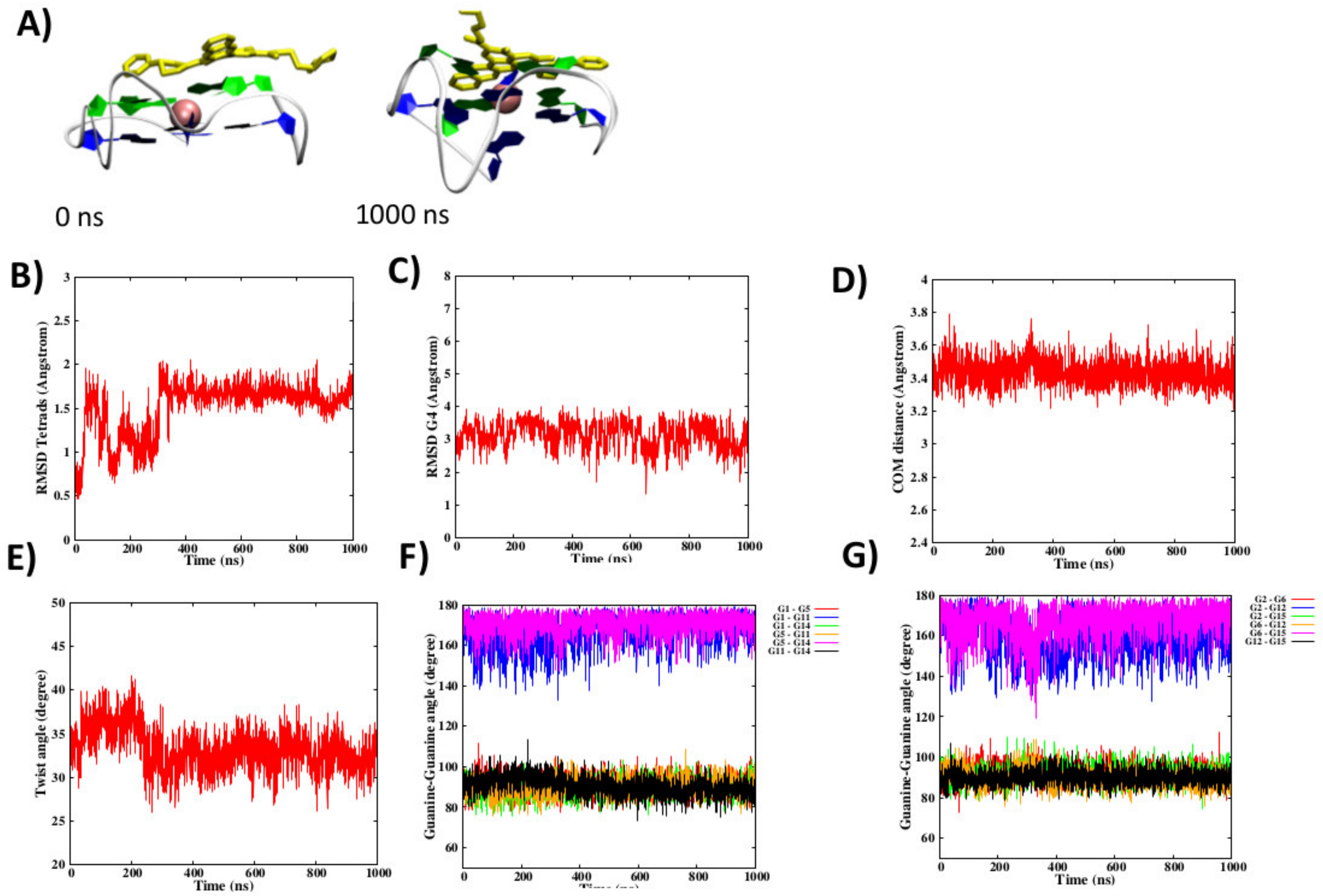


**Figure S11:** Initial and final snapshot of the first MD simulations of the RG-1 sequence interacting with QRF in the top binding pose (A). RMSD of the guanine in the tetrad (B) and of the flexible loop (C), distance between the tetrad (D), twist Ω angle (E), and ϕ, ϕ' angles for the first (F) and second (G) tetrad.

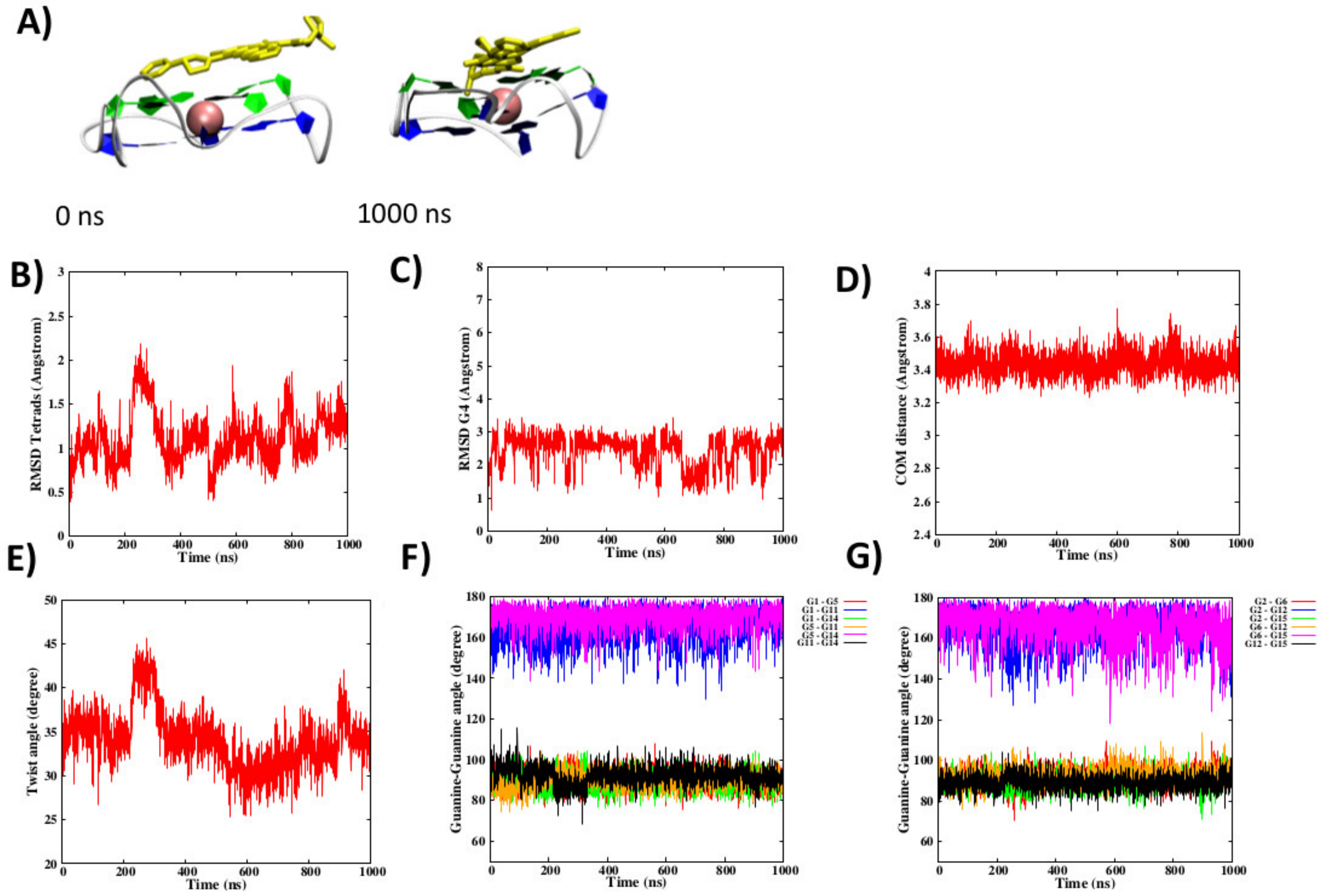


**Figure S12:** Initial and final snapshot of the second MD simulations of the RG-1 sequence interacting with QRF in the top binding pose (A). RMSD of the guanine in the tetrad (B) and of the flexible loop (C), distance between the tetrad (D), twist $\Omega$ angle (E), and $\phi$, $\phi'$ angles for the first (F) and second (G) tetrad.

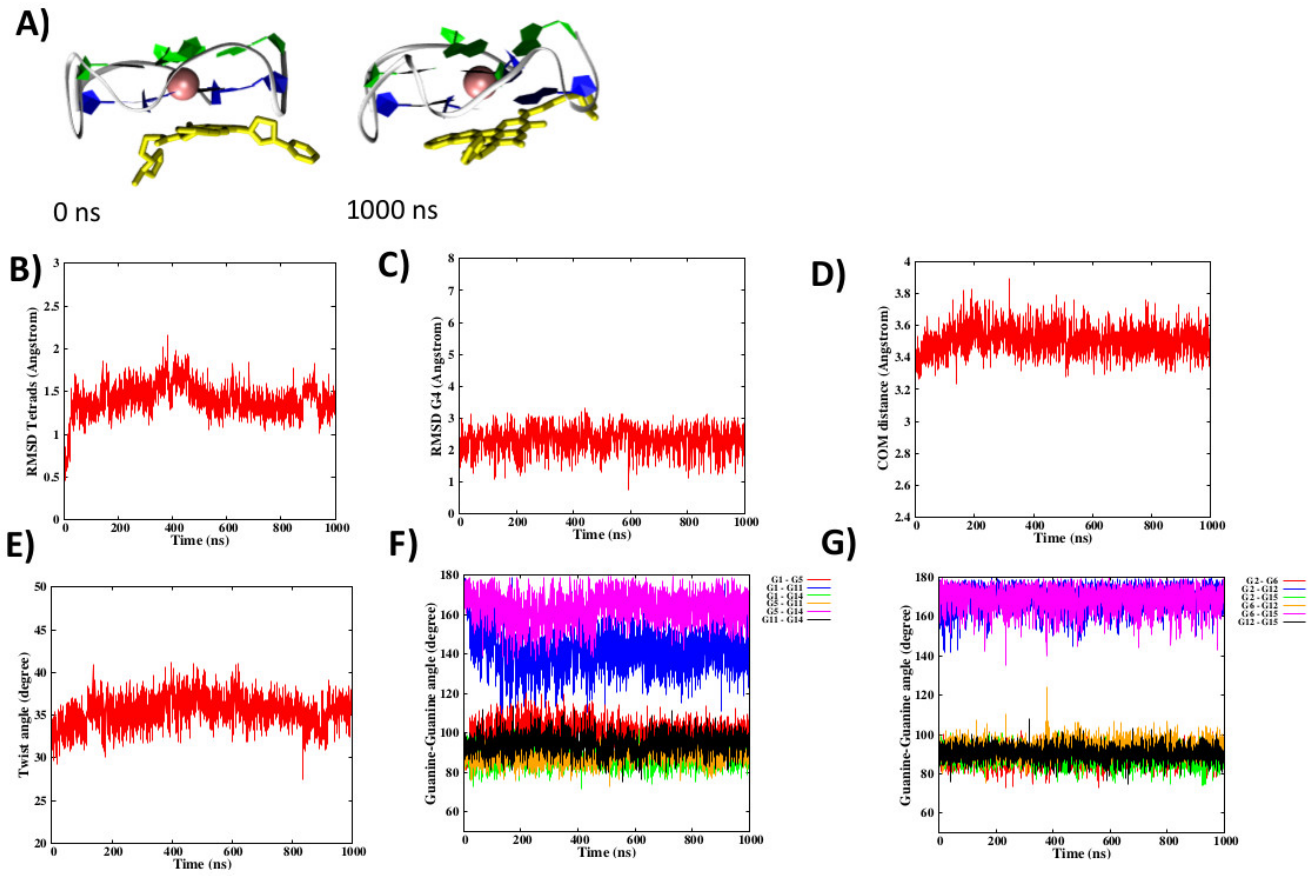


**Figure S13:** Initial and final snapshot of the first MD simulations of the RG-1 sequence interacting with QRF in the down binding pose (A). RMSD of the guanine in the tetrad (B) and of the flexible loop (C), distance between the tetrad (D), twist Ω angle (E), and ϕ, ϕ' angles for the first (F) and second (G) tetrad.

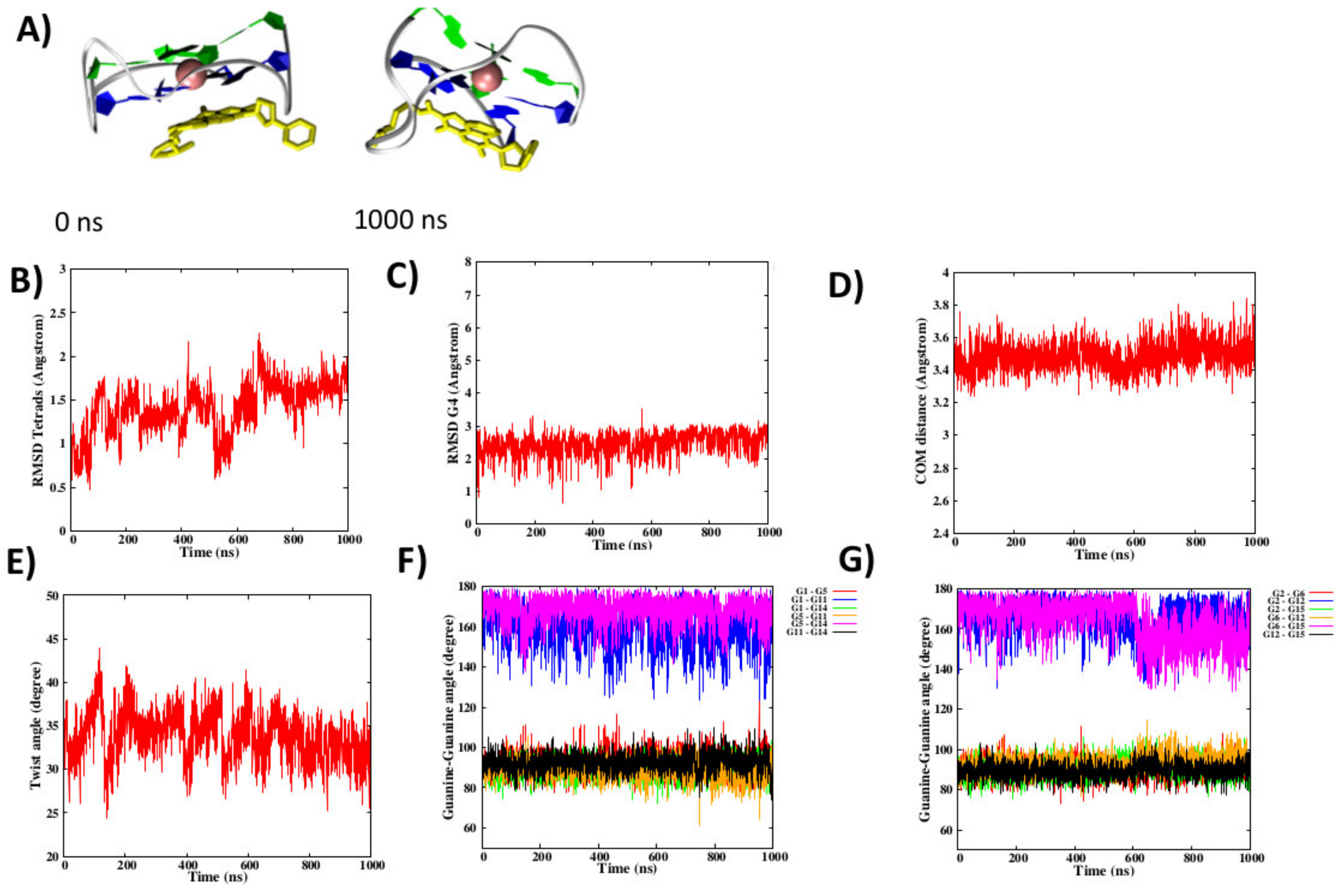


**Figure S14:** Initial and final snapshot of the second MD simulations of the RG-1 sequence interacting with QRF in the down binding pose (A). RMSD of the guanine in the tetrad (B) and of the flexible loop (C), distance between the tetrad (D), twist $\Omega$ angle (E), and $\phi$, $\phi'$ angles for the first (F) and second (G) tetrad.
